# Minimizing the influence of magnetic vestibular stimulation inside MRI-scanners by adjusting head position

**DOI:** 10.1101/2025.08.22.670281

**Authors:** Laura Schaffeld, Leonie Behle, Axel Lindner, Hans-Otto Karnath

## Abstract

Magnetic resonance imaging (MRI) offers detailed diagnostic insights but also unintentionally induces vestibular side effects by its static magnetic field. Magnetic vestibular stimulation (MVS) may cause dizziness and confound behavioral and physiological measures in fMRI by introducing an unavoidable mixture of neural activation in vestibular projection areas, driven both by the signal of interest and by vestibular system activation. This study investigated how different head orientations within a 3T MRI scanner influence the MVS-induced vestibulo-ocular reflex (VOR) in complete darkness, with the goal of identifying a head position that minimizes or eliminates MVS effects. Results revealed a linear relationship between head pitch and the horizontal VOR (Experiment 1), as well as between head roll and the vertical VOR (Experiment 2). Across participants, the horizontal VOR was eliminated at an average forward head pitch of 24.4°, while the vertical VOR was nullified at an average roll angle of 15.9° towards the right shoulder. A head position combining this forward pitch with this rightward roll effectively minimized both horizontal and vertical VOR components across subjects (Experiment 3). The findings provide a practical solution to reduce the impact of MVS effects in MRI with important implications for improving data quality in neuroscientific fMRI studies and patient comfort during clinical imaging.

## 1. Introduction

The vestibular system of the inner ear has a fundamental function in detecting head motion and spatial orientation, ensuring stable visual perception and coordinated motor responses. By integrating signals from the vestibular labyrinth with proprioceptive and visual information, it supports postural control and gaze stabilization (Angelaki & Cullen, 2008). Interestingly, the vestibular system cannot only be stimulated by self-motion but also by exposure to strong magnetic fields, such as those permanently present in magnetic resonance imaging (MRI) scanners. This phenomenon, known as *magnetic vestibular stimulation* (MVS), has been discovered based on observations by Marcelli et al. (2009) and has since been reliably replicated across scanners of various field strengths (see Ward et al., 2019, for review). The underlying mechanism derives from an interaction between the scanner’s static magnetic field and ionic currents in the endolymphatic fluid above the utricle. This interaction acts primarily on the cupulae of the lateral and anterior semicircular canals (Roberts et al., 2011; Otero-Millan et al., 2017; Arán-Tapia et al., 2025) and predominantly induces a horizontal vestibular ocular reflex (VOR), akin to that triggered by actual head rotation (Otero-Millan et al., 2017).

Importantly, the effects of MVS extend beyond the aforementioned reflexive oculomotor responses. Recent work has demonstrated that MVS also alters spatial attention and exploration. Lindner et al. (2021) showed that exposure to a 3 T magnetic field induces a sustained horizontal bias in overt attention during a visual search task conducted in darkness. The attentional shift induced by MVS remains consistently present for the entire time the subject is in the scanner (Smaczny et al., 2024), as does the MVS-induced VOR (Jareonsettasin et al., 2016; Go et al., 2022; Smaczny et al., 2024). Besides the aforementioned attentional and behavioral biases induced by MVS, Boegle and coworkers (2016) further reported fluctuations in resting state fMRI activity varying with different static magnetic field strengths (1.5 T vs. 3 T; also compare Roberts et al., 2011). These modulations affected brain regions related to vestibular and oculomotor functions.

These aforementioned findings challenge the common interpretation of behavioral and BOLD-activity measures obtained in fMRI experiments. It indicates that during the entire fMRI acquisition time the neuronal activation in the (cortical) projection areas of the vestibular system represent an unresolvable mixture of the signal of interest and the signal evoked by the activation of the vestibular system. One could argue that in experimental settings without absolute darkness the VOR can most likely be supressed. Yet, even when fixation of a visual stimulus allows suppression of the VOR the vestibular stimulation pertains and could influence cognition and behavior: Buettner and Büttner (1979) elegantly demonstrated that during VOR suppression there is little change in the maximal firing rate of single neurons recorded in the vestibular nuclei of monkeys. The activity in 80% of the neurons was reduced by only less than 10%.

For that reason, one should thus seek for MRI-settings that might help abolishing or minimizing this impact on behavioral and BOLD-activity measures obtained in fMRI experiments. The known dependency of the MVS-induced horizontal VOR on head pitch position seems helpful for this: Pitching the head towards subjects’ chest reduces the horizontal (and torsional) VOR (Roberts et al. 2011; Boegle et al., 2016). However, also the vertical domain should be considered. This is because it has been consistently reported that simply lying in the supine position in darkness (without any MRI) does cause an upbeat nystagmus (and thus a downward VOR) (Bisdorff et al., 2000; also compare Young et al., 2020). Also in our own previous work, we did document such a downward bias of the VOR in subjects lying in the supine position on the scanner table and this bias was present both inside and outside the MRI (Lindner et al., 2021; Smazcny et al., 2024). Preliminary observations by various groups (Roberts et al., 2011; Go et al., 2022) as well as a theoretical model (Arán-Tapia et al, 2025) suggest that “rolling” a subject’s head towards the shoulder inside an MRI scanner modulates the vertical VOR.

Based on these considerations and previous findings, we here tried to empirically derive the head pitch and/or head roll position in individuals, for which the horizontal and vertical effects of MVS (as quantified through the resulting VOR) inside an MRI scanner is nulled. In addition, we wanted to see whether combining these head pitch and roll positions in an independent experiment would effectively abolish both horizontal and vertical VOR at the same time.

## 2 Methods

### 2.1 Participants

To investigate the influence of *head pitch* (relative to the b_0_ magnetic field-vector) on the VOR, a total of thirteen healthy subjects participated in *Experiment 1*. The eye-tracking data of three subjects were excluded from the analysis, either due to a significant deviation of eye gaze which did not allow for pupil tracking or due to continuous tracking-artifacts introduced by eyelashes. Therefore, data from 10 subjects (3 males, mean age = 26.3 years, SD = 2.2 years) were included in the analysis.

In *Experiment 2*, which studied the VOR as a function of *head roll* (relative to the b_0_-vector) by tilting the head towards the shoulder, an additional 12 healthy subjects participated (5 males, mean age = 26.1, SD = 4.0). All participants either had normal vision or used MRI-compatible glasses for correction.

Based on the results obtained from the first two experiments, we conducted an additional, third experiment. In this *Experiment 3*, we aimed to experimentally validate whether a combination of head pitch and roll (values according to the findings from Experiments 1 and 2) could null the VOR in both the horizontal and vertical domain. To determine the required sample size to provide enough power in support of this null-hypothesis, we conducted a respective power analysis using G*Power while building on an effect size derived from Smaczny and colleagues (2024; effect size: *dz* = 1.49, *α* = 0.05, and power [1 – β] = 0.95). The analysis indicated that a minimum of 7 subjects would be sufficient to probe for the presence/absence of an effect (one-tailed paired-samples t-test). Accordingly, we recruited 7 additional healthy participants (5 males; mean age = 29.4, SD = 1.3)

None of the participants reported a history of vestibular or neurological disease. The experiments were approved by the local ethics board and have been performed in accordance with the revised Declaration of Helsinki. Prior to the experiment, all volunteers provided informed consent according to the guidelines of our institutional ethics committee.

### 2.2 Apparatus and Stimuli

In all three experiments, a 3 T Siemens MAGNETOM Prisma MRI scanner was employed. Since no imaging was conducted, neither radiofrequency nor gradient fields were applied, and only the static magnetic field was present. Participants always entered the MRI scanner in the standard head-first orientation, with the static magnetic field vector (b_0_-vector) pointing from subjects’ toes to their head.

For stimulus generation and control, a Windows 10 Laptop PC running Matlab R2015b (The MathWorks Inc.) with the Cogent2000 Cogent Graphics Toolbox (FIL, ICN, LON at the Wellcome Department of Imaging Neuroscience, University College London) was utilized. Visual stimuli for calibrating the eye-tracker were presented through a VisuaStim Digital MRI-compatible fMRI system with video goggles (Resonance Technology, Inc.; 800 × 600 resolution and 60 Hz refresh rate; 30° x 24° field of view). The use of video goggles allowed reliable presentation of visual calibration stimuli even when participants’ head positions were varied during the study. Importantly, the goggles’ display could be turned off completely, which enabled us to assess participants’ VOR in complete darkness. 2D-eye movements were recorded using the analog fMRI Eyetracker Camera (Resonance Technology, Inc.; sampling rate: 60Hz; camera resolution: 320×240 pixels), which is integrated into the VisuaStim video goggle system (compare Figure 1). The camera captured the participant’s right eye via a semi-transparent mirror in front of the display. The video signal was digitized using the ViewPoint monocular integrator system (Arrington Research) and eye position was monitored via dark pupil tracking implemented in the ViewPoint software (Arrington Research, version 2.8.3.437). The system was operated on a secondary Windows 10 PC, remotely controlled by the ViewPoint Ethernet client running on the control PC.

**Figure 1.**
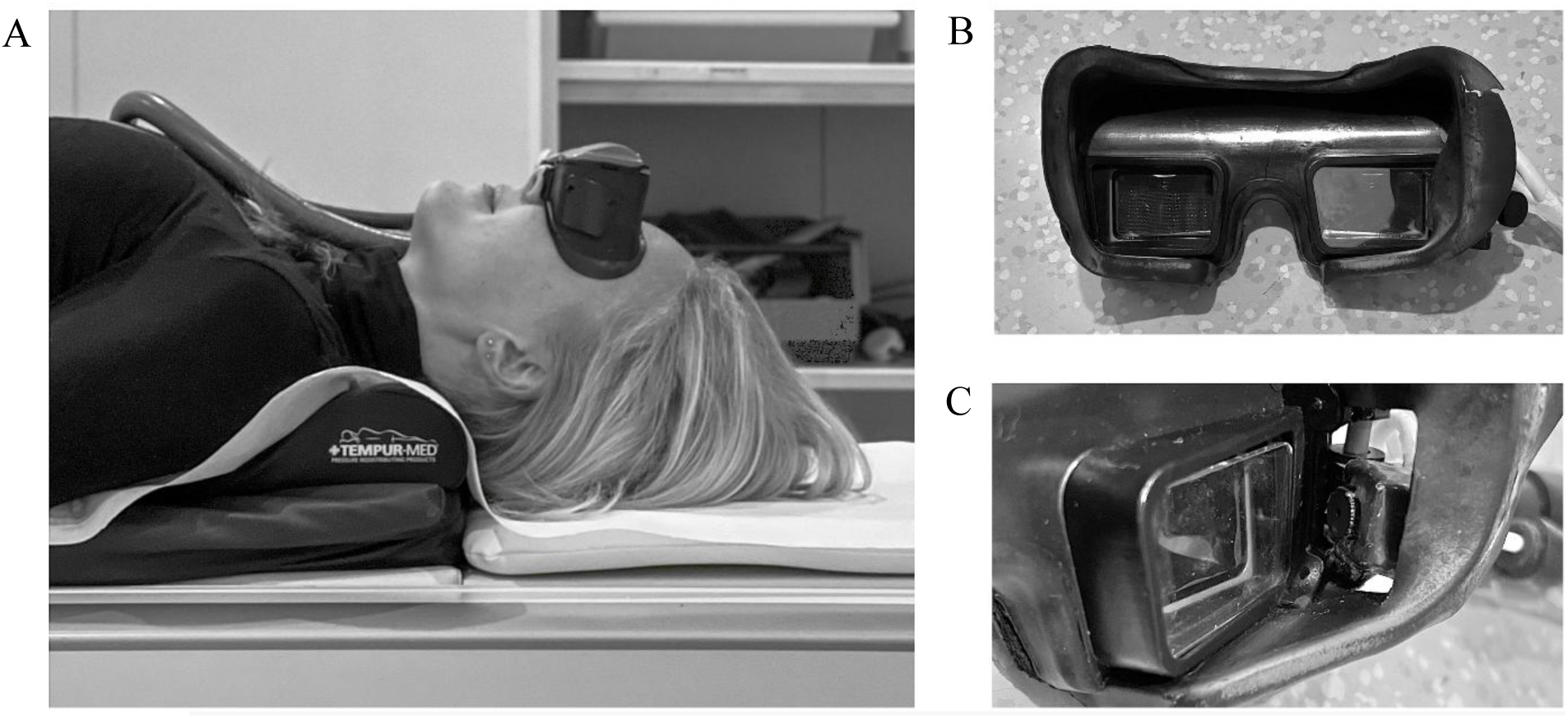
Experimental setup with MRI-compatible eye-tracking goggles. (A) Participants were wearing the VisuaStim Video Goggle system for stimulus presentation and to record eye movements using the integrated analogue eye-tracking video camera. The use of the goggles ensured consistent visual stimulus presentation despite varying head positions. (B) Interior view of MRI-compatible goggles with integrated display for visual stimulus presentation. (C) Close-up of infrared camera within goggles capturing right eye movement through a semi-transparent mirror in front of the right display.

### 2.3 Experimental procedure

To investigate the effects of a magnetic field on the VOR, two studies were conducted. The first study focused on the influence of head pitch inside the MRI scanner bore on the VOR, while the second study targeted the effect of head roll on the VOR. Both experiments consisted of two tasks: eye calibration and nystagmus assessment performed in complete darkness to measure horizontal and vertical eye movement.

#### Experiment 1

In the first experiment, three head pitch positions were used to modulate the effect of MVS on the VOR (Figure 2A). One position involved a backward tilt of the head of -30° relative to the vertical axis and the canthus-tragus line, maximizing the MVS effect on the horizontal semicircular canals as described by Roberts et al. (2011) and Boegle et al. (2016). Another position (0°) represented a head orientation, with no tilt relative to the vertical axis thereby mimicking the typical head positioning during MRI. Finally, we employed a forward tilt of +30°, which should reduce the influence of MVS on the horizontal semicircular canals (cf. Roberts et al., 2011; Boegle et al., 2016). The sequence of head pitch positions was counterbalanced among participants to rule out potential order effects. Cushions and pillows were used to achieve the desired head pitch angles while ensuring participant comfort.

**Figure 2.**
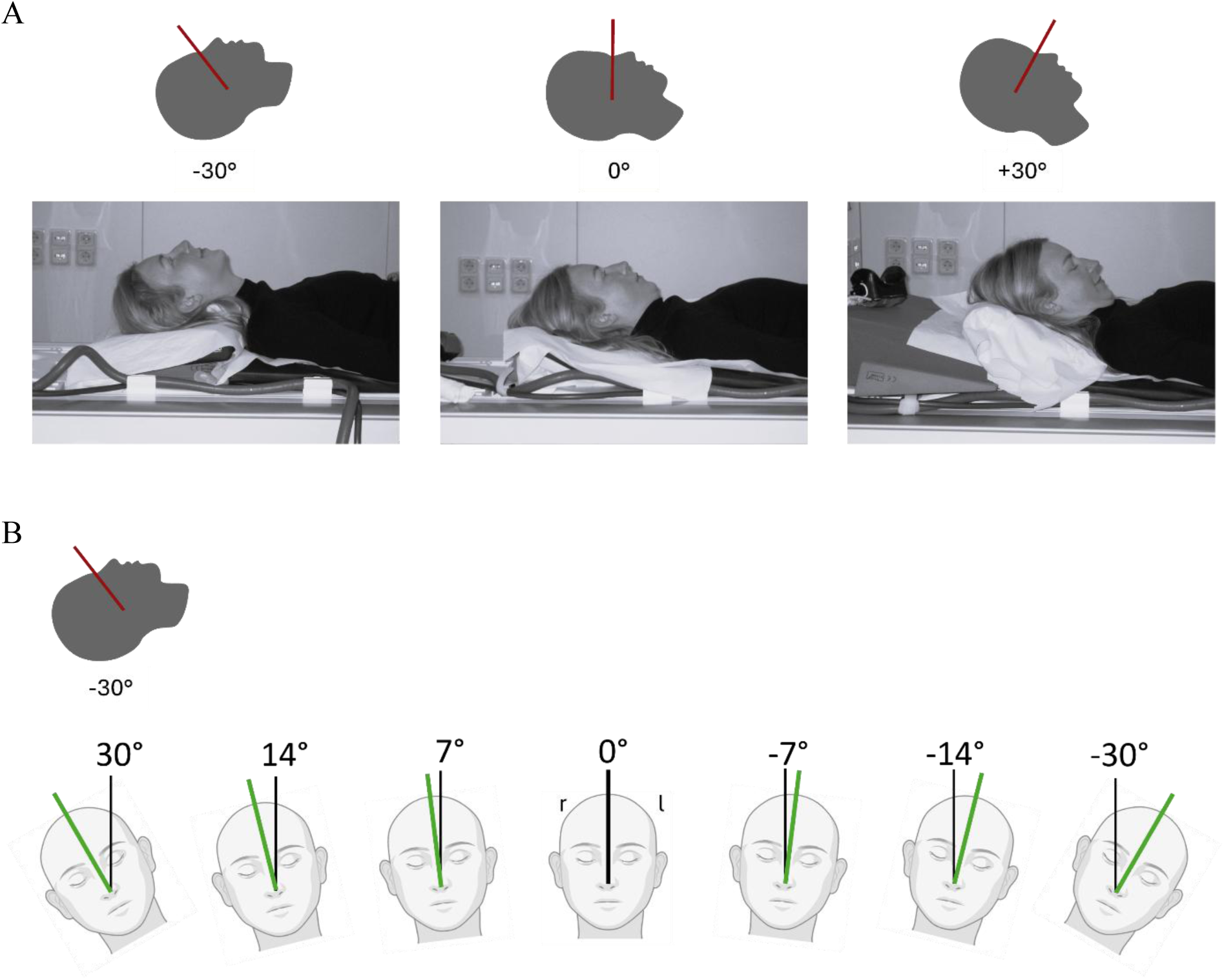
Head positions used in the Experiments 1 and 2. The red line represents the canthus-tragus line. (A) For Experiment 1, three head pitch positions were tested: -30°, 0°, and +30° relative to the vertical axis and the canthus-tragus line. (B) For Experiment 2, participants maintained a head pitch of -30° while completing tasks across seven head roll positions: -30°, -14°, -7°, 0°, 7°, 14°, and 30°. Roll angles were measured relative to two reference lines: the green line indicates the nasal bridge, and the black line represents the sagittal body axis. Negative values indicate tilt towards the participant’s left shoulder, positive values tilt towards the right.

In all three head pitch positions of the first experiment, participants first completed an *eye calibration task* to ensure proper alignment with the subsequent *nystagmus assessment*, as adjustments to head position might affect the goggles’ calibration. The calibration involved visual fixation on a central cross, followed by four peripheral visual crosses presented sequentially on a black background at predefined coordinates (0°/0°, 0°/4°, 5.6°/0°, 0°/-4°, and -5.6°/0° visual angle) for three seconds each. This sequence was repeated twice for accuracy. Participants then were instructed to maintain their gaze straight ahead in complete darkness for one minute, while eye movements were recorded to assess possible nystagmus. During this period, the goggles’ displays were switched off and eye movement recordings were analyzed to derive slow-phase velocities as a measure of the VOR.

#### Experiment 2

The second experiment assessed the VOR by systematically varying the roll angle of the head while maintaining a constant head pitch of -30° relative to the vertical axis (Figure 2B). Seven roll positions were tested, ranging from -30° to +30° (-30°, -14°, -7°, 0°, 7°, 14°, and 30°). Negative roll values corresponded to tilts towards the participant’s left shoulder, while positive values corresponded to tilts towards the right shoulder. To ensure stability in the tilted positions, the head was supported on both sides with sandbags. Participants were initially positioned at 0° head roll outside the MRI scanner to establish an outside control condition. This outside control was always followed by the 0° head roll-condition inside the scanner, namely as a manipulation check to probe for the presence of the expected MVS-induced horizontal VOR (cf. Roberts et al., 2011). The remaining head roll positions were tested in a pseudorandomized order to mitigate time-related effects. As compared to the first experiment, the order of the tasks was reversed. The one-minute *nystagmus assessment* was performed first, followed by the *eye calibration task* in each roll position.

#### Experiment 3

Based on the results obtained from the first two experiments (see results section), we conducted an additional, third experiment in which we combined at the same time the head pitch position that nulled the MVS-induced horizontal VOR (Experiment 1) and that head roll position that nulled the MVS-induced vertical VOR (Experiment 2). With tilt values rounded to 5° steps, a combined head tilt condition in which participants’ heads were positioned at 25° forward pitch and 15° roll toward the right shoulder was used. Each participant was tested in three head positions: this combined tilt (25° pitch, 15° roll) position, the standard (0° pitch, 0° roll) supine position, and the backward tilt (-30° pitch, 0° roll) position, in which we expect the MVS-induced VOR to be maximal. We used this latter position as a manipulation check for MVS, since the magnetic field effect is strongest in this orientation and a VOR is reliably expected. To rule out that habituation effects could explain the absence of a VOR in the combined tilt condition, it was always presented as a second inside MRI-run while the standard supine and the backward tilt conditions were assigned to the first and third inside MRI-run in a counterbalanced fashion. Hence, if a VOR was still detectable in these latter two conditions, habituation could not account for an absence of the VOR in the combined tilt condition. Head positions were maintained using a combination of the MRI head coil and supporting cushions. As in the first two experiments, eye calibration and nystagmus assessments were again conducted for each position inside the MRI scanner.

### 2.4 Eye Movement Analysis

Eye movement data were processed offline using a consistent analysis pipeline across both experiments (custom MATLAB R2017B scripts (MathWorks)) to ensure comparability of results. A five-point calibration procedure was performed separately for each head position. Next, calibration was applied to the data from the *nystagmus assessment*. The calibrated 2D eye data were then filtered with a 10 Hz second-order digital low-pass filter and saccades were identified using an absolute eye velocity threshold of 15°/s. Eye velocity was calculated via two-point differentiation of the position data, with saccades marked from the first sample exceeding the threshold to the first sample below. Horizontal and vertical eye velocity traces were desaccaded by treating saccade intervals as missing data points. Blink artifacts were also excluded. Individual horizontal and vertical slow phase VOR were derived from these velocity traces.

### 2.5 Statistical Analysis

A significance level of α = 0.05 was used for all statistical tests. All analyses were conducted using IBM SPSS Statistics (Version 29.0.2.0). For the first two experiments, the median horizontal and vertical slow phase velocity from the nystagmus assessment was calculated for each participant at each head position. The respective data were first tested for normality using a Shapiro-Wilk test. In case of a normal distribution, the influence of head position on slow phase VOR was tested with a repeated-measures ANOVA (rmANOVA). For the rmANOVAs, the Mauchly test for sphericity was conducted in all cases. If sphericity was violated, the Greenhouse-Geisser correction was applied. Statistical contrasts provided by the rmANOVA were additionally examined for both “linear” and “quadratic” effects. In cases where the normality test indicated a non-normal distribution, we used a non-parametric equivalent for the rmANOVA, namely the Friedman test. Additionally, rmANOVAs were used to probe for any influence of measurement order, e.g. reflecting whether the VOR adapted the more time subjects spent inside the MRI (cf. Jareonsettasin et al., 2016; Smaczny et al., 2024). In Experiment 2, the measurement protocol inside the MRI always started with the 0° roll condition as a manipulation check (compare below). Thus, the horizontal VOR in this position should have been affected the least by adaptation (compare supplementary Figure S1). Instead, all other roll positions were presented only later and in a counterbalanced order. For investigating the influence of head roll position on the horizontal VOR the 0° position will therefore not be considered.

In Experiment 1, one-tailed one-sample t-tests were conducted additionally, namely to control for biases of the vertical VOR in the supine position (cf. Bisdorff et al., 2000; also compare Young et al., 2020). If normality was rejected, a one-tailed Wilcoxon signed-rank test served as a non-parametric alternative. In Experiment 2, the VOR values measured at the 0°-head roll position both outside (i) and inside (ii) the MRI scanner were compared using one-tailed paired t-tests. This served as a manipulation check to probe for the presence of a horizontal VOR induced by MVS (cf. Roberts et al., 2011).

Finally, in Experiments 1 and 2 regression analyses were performed on the slow phase velocity data in each individual as a function of head position, focusing on the respective x-intercepts, y-intercepts, correlation coefficients (r), and slopes. The x-intercept was analyzed to determine the head position of each participant at which no VOR was present, while the y-intercept provided information about the baseline VOR in the standard head position within the MRI. The correlation coefficient (r) was used to assess the goodness of fit for the linear model in individuals, and the slope quantified the sensitivity of the horizontal and vertical VOR to varying head pitch and head roll position, respectively. The resulting individual correlation coefficients, the x-intercepts, and the slopes of the regressions were further analyzed on the group level. To this end, we first assessed whether they are normally distributed using the Shapiro-Wilk test. If normality was confirmed, one-sample t-tests were conducted to test whether these regression parameters significantly differed from 0. If normality was rejected, Wilcoxon signed-rank tests were applied as a non-parametric alternative.

Lastly, to evaluate the absence of MVS effects in Experiment 3 we used Bayes-Factor (BF) analyses, following the approach described by Dienes (2011; see Figure A4 of that paper). This method is particularly appropriate, as it allows us to directly quantify evidence for the presence of a VOR (H₁; BF>3) versus its absence (H₀; BF<1/3). Unlike frequentist statistical approaches, which depend heavily on sample size to yield reliable p-values and power, Bayesian methods assess the strength of evidence for competing hypotheses irrespective of sample size (Dienes, 2011; Wagenmakers et al., 2018). For our two-tailed BF analyses, population values from a study by Smaczny et al. (2024) were used as priors. Specifically, the mean horizontal (1.64°/s) and vertical VOR (–0.74°/s) were entered as estimates for the priors’ standard deviation while assuming means of 0°/s (compare Appendix and Fig. A4 in Dienes, 2011).

## 3 Results

### 3.1 Experiment 1: Influence of head pitch position inside a 3 T MRI scanner

Figure S1 (cf. supplement) illustrates the eye movement data from one representative participant and Figure 3 the distribution of the median slow phase velocity for each participant across the different head pitch positions (-30°, 0°, +30°) while lying inside the 3 T scanner. When the head was pitched approximately -30° backward, all participants exhibited a rightward slow phase of nystagmus, with an average horizontal velocity of 1.63 °/s (*SE* = 0.28 °/s). In the “neutral position” (0°), all except one participant showed a reduction in their rightward slow phase, in one of whom there was even a reversal towards the left. Mean velocity at 0° pitch angle was 0.66 °/s (*SE* = 0.21 °/s). This leftward change in velocity continued for the +30° pitch angle, where the horizontal slow phase reached an average of -0.123 °/s (*SE* = 0.20 °/s).

**Figure 3.**
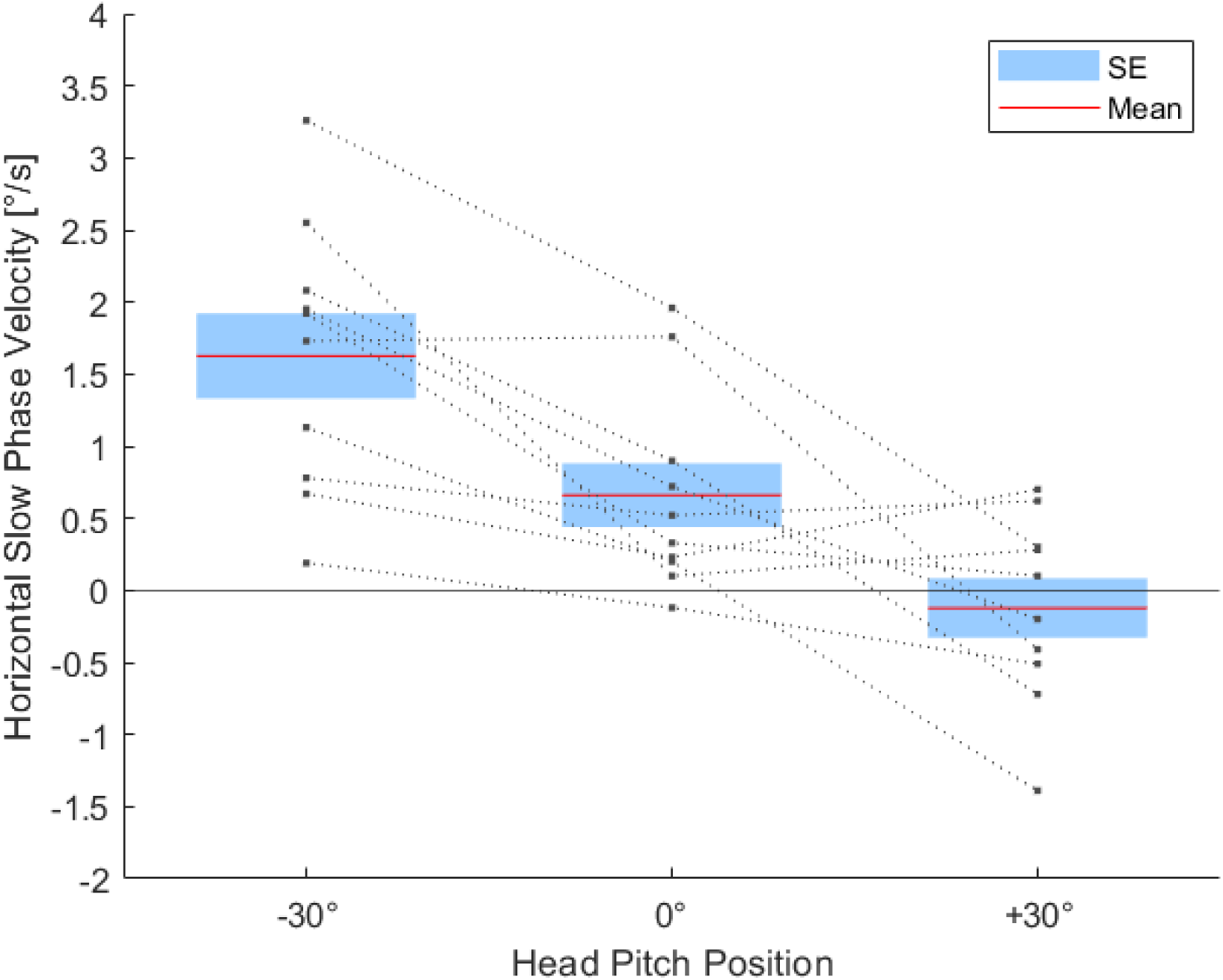
Horizontal slow phase velocity [°/s] for each head pitch position across all subjects. Dotted lines represent the desaccaded median horizontal eye velocity of individual subjects (circles, N = 10) for the three head positions -30°, 0° and +30°. Positive values indicate a rightwards slow phase, negative values signify a VOR towards the left. The red lines and blue boxes reflect the mean slow phase velocity and the standard error (SE) across all subjects.

An rmANOVA was conducted to statistically evaluate the obvious relationship between head pitch position and horizontal slow phase, revealing a highly significant effect (*F*(2, 18) = 17.74, *p* < 0.001, *η²p* = 0.66). This confirms earlier conclusion that head pitch significantly influences the MVS-induced horizontal VOR (Roberts et al., 2011; Boegle et al., 2016). A repeated-measures contrast analysis further indicated a significant “linear effect” of head pitch position on the horizontal VOR (*F*(1, 9) = 26.11, *p* < 0.001, η²p = 0.74), while no “quadratic effect” could be revealed (*F*(1, 9) = 0.20, *p* = 0.666, *η²p* = 0.02). To further investigate this relationship, a linear regression analysis was performed in each individual. A one-tailed one-sample *t*-test confirmed that the respective slopes of the individual regressions were significantly smaller than zero (*t*(9) = -5.11, *p* < 0.001). As the Shapiro-Wilk test indicated that the respective correlation coefficients (*r*) of these regressions were not normally distributed (*p* < 0.001), a two-tailed Wilcoxon signed-rank test was conducted, confirming that *r* was significantly different from zero (*W* = 55, *p* = 0.005). These findings demonstrate a progressive decrease in MVS-induced horizontal slow phase velocity with increasing head pitch towards the chest. The median x-intercept, i.e. the pitch position for which the participants showed no horizontal VOR, was 24.4° (*IRQ* = 34.4°), and the mean y-intercept and thus the average baseline activity of the participants in the standard head position was 0.18 °/s (*SE* = 0.72 °/s).

The same analyses were then conducted for vertical VOR. Figure 4 illustrates the distribution of the vertical slow phase velocity for all individual subjects in each head pitch position. Notably, the vertical VOR exhibited a consistent negative velocity for all subjects and across all head pitch positions, indicating a downward bias. The mean velocity of the slow phases for position -30° pitch angle was -0.93 °/s (*SE* = 0.20), -0.82 °/s (*SE* = 0.20) for 0°, and -1.05 °/s (*SE* = 0.22) for +30°.

**Figure 4.**
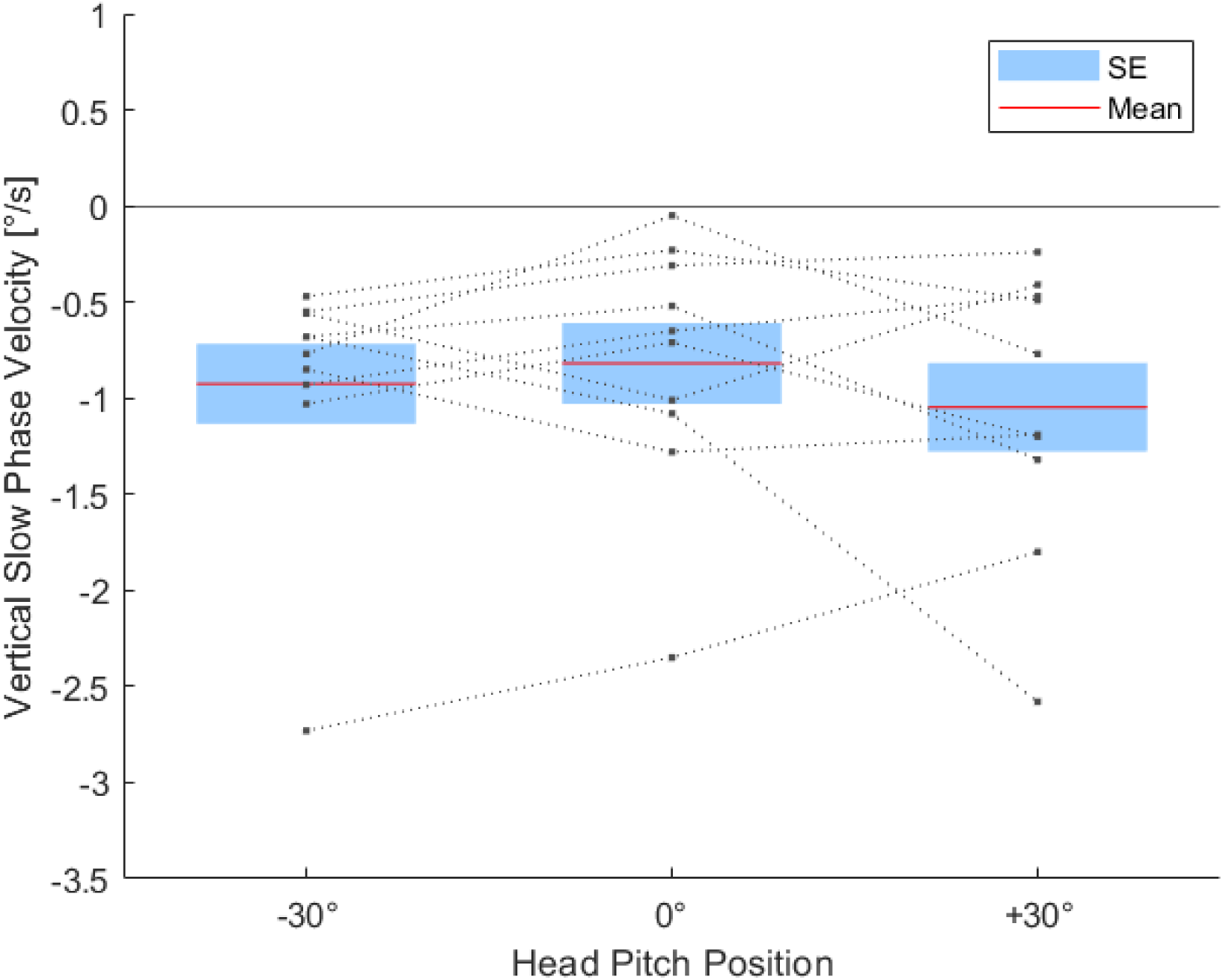
Vertical slow phase velocity [°/s] for each head pitch position across all subjects. Dotted lines represent the desaccaded horizontal eye velocity of individual subjects (circles, N = 10) for the three head positions -30°, 0° and +30°. Positive values indicate a rightwards slow phase, negative values signify a VOR towards the left. The red lines and blue boxes reflect the mean slow phase velocity and the standard error (SE) across all subjects.

To evaluate the relationship between head pitch position and the vertical VOR we performed a Friedman test as a Shapiro-Wilk test indicated non-normality for position -30° pitch angle (*p* < 0.001). The Friedman test revealed no significant differences in the vertical VOR between different head pitch positions (*χ²*(2) = 0.67, *p* = 0.717). These results are in line with previous findings demonstrating that vertical VOR inside MRI-scanners remains largely unaffected by changes in head pitch (Otero-Millan et al., 2017; Lindner et al., 2021; Arán-Tapia et al., 2025). Nevertheless, all three positions revealed a significant downward VOR (two-tailed Wilcoxon signed-rank test for -30° pitch angle: *W* = 0, *p* = 0.005; one-tailed one-sample t-test for 0°: *t*(9) = -3.89, *p* = 0.002 and for +30°: *t*(9) = -4.53, *p* < 0.001).

### 3.2 Experiment 2: Influence of head roll angle inside a 3 T MRI scanner

In the supine position, a downward VOR in darkness was previously reported in normal laboratory settings without MRI (Bisdorff et al., 2000; also compare Young et al., 2020) and was also consistently present in Experiment 1 (cf. Fig. 4). In Experiment 2, we first replicated the downward VOR outside the scanner in the neutral head roll position at 0°. The mean vertical slow phase velocity amounted to -0.64 °/s (*SE* = 0.30 °/s) and was significantly smaller than 0°/s (one-tailed one-sample t-test: *t*(11) = -2.18, *p* = 0.026). We next compared whether entering subjects into the scanner would modulate this effect. The vertical slow phase velocity inside the scanner was now 0.12 °/s (*SE* = 0.44 °/s). The velocities at head position 0° thereby turned out to be significantly different between outside and inside (*t*(11) = 2.81, *p* = 0.017). Accordingly, a one-tailed one-sample t-test, investigating whether the slow phase velocity differs from zero showed no significance inside the scanner (*t*(11) = 0.26, *p* = 0.398).

Figure S2 (cf. supplement) illustrates the eye movement data from one representative participant and Figure 5 the distribution of the vertical slow phase velocity across all seven head positions measured inside the scanner. The mean vertical velocities for each roll position were -0.93 °/s at position -30° to the subject’s left (*SE* = 0.26 °/s), -0.79 °/s at position -14° (*SE* = 0.28 °/s), -0.58 °/s at position -7° (*SE* = 0.34), 0.12 °/s at usual head position of 0° (*SE* = 0.44 °/s), 0.04 °/s at position 7° to the subject’s right (*SE* = 0.39 °/s), 0.09 °/s at position 14° (*SE* = 0.33 °/s), and 0.26 °/s at position 30° (*SE* = 0.26 °/s). The general trend suggests a progressive increase in vertical slow phase magnitude with increasing head tilt from left to right, which aligns with a previous experiment by Go et al. (2022).

**Figure 5.**
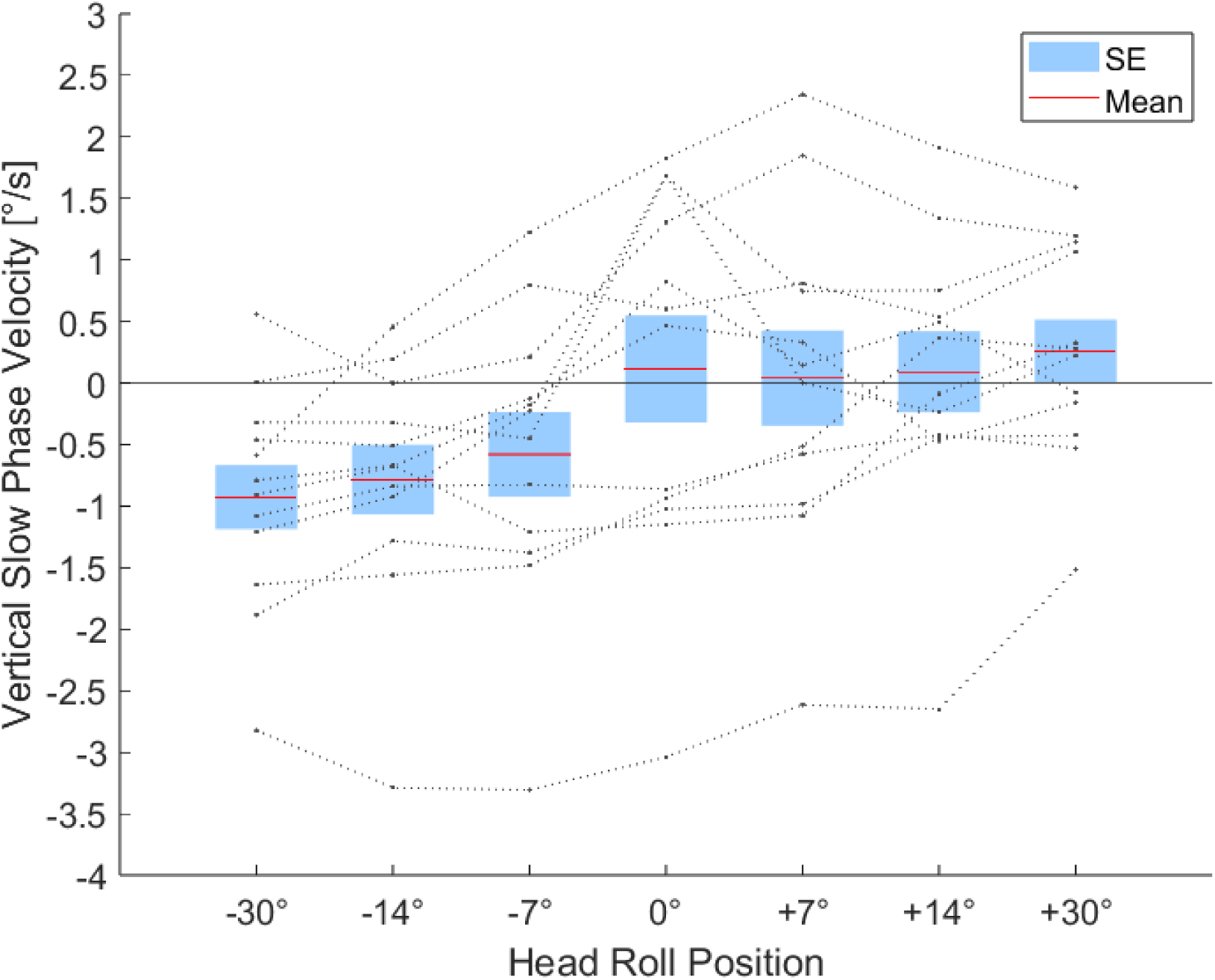
Vertical slow phase velocity [°/s] for each head roll position across all subjects. Dotted lines represent individual subject data (circles, N = 12) for the seven head positions. Negative values indicate downward slow phase, positive values indicate an upward direction. The red lines and blue boxes reflect the mean slow phase velocity and standard error (SE) across all subjects.

To assess the effect of head roll angle on vertical VOR, an rmANOVA was conducted, and the main effect of the roll position was highly significant (*F*(6, 66) = 12.35, *p* < 0.001, *η²p* = 0.53). In addition, contrast analysis revealed a significant linear (*F*(1, 11) = 55.63, *p* < 0.001, *η²p* = 0.84, but no quadratic effect (*F*(1, 11) = 1.21, *p* = 0.295, *η²p* = 0.10). To further investigate the linear relationship, regression analysis was performed. A one-tailed one-sample t-test confirmed that the slope (*t*(11) = 7.58, *p* < 0.001) and the correlation coefficient r (*t*(11) = 12.48, *p* < 0.001) significantly differed from zero, indicating a systematic linear change in vertical VOR depending on the head tilt position. On average the participants had an x-intercept and therefore no vertical VOR at 15.9° (*SE* = 13.6°), while the average baseline activity of the participants in the standard head position at 0° was -0.25 °/s (*SE* = 0.31 °/s). Both measures were not significantly different from zero, as revealed by a one-tailed one-sample t-test (x-intercept: *t*(11) = -0.829, *p* = 0.212; y-intercept: *t*(11) = 1.175, *p* = 0.132).

The same analysis was conducted for horizontal VOR across all head roll positions. First, however, we performed a manipulation check to assess whether the expected MVS-induced horizontal VOR was present in the neutral 0° head roll position: Comparing the horizontal slow phase velocity outside the MRI scanner (*mean* = 0.05 °/s, *SE* = 0.15 °/s) and inside (*mean* = 3.66 °/s, *SE* = 0.67 °/s), a one-tailed one-sample *t*-test confirmed that horizontal slow phase significantly differed from zero inside the scanner (*t*(11) = 5.50, *p* < 0.001) but not outside (*t*(11) = 0.32, *p* = 0.378). A paired-samples *t*-test further revealed a significant difference between these conditions (*t*(11) = 5.24, *p* < 0.001), indicating that the magnetic field induced the expected horizontal VOR.

Figure 6 depicts the distribution of the median horizontal slow phase velocity across participants for the additional roll positions inside the scanner. As expected, these mean horizontal velocities hardly varied and were 1.80 °/s at position -30° (*SE* = 0.47 °/s), 1.74 °/s at position -14° (*SE* = 0.25 °/s), 1.77 °/s at position -7° (*SE* = 0.33 °/s), 1.94 °/s at position 7° (*SE* = 0.30 °/s), 1.83 °/s at position 14° (*SE* = 0.32 °/s), and 1.73 °/s at position 30° (*SE* = 0.26 °/s). Accordingly, a respective rmANOVA revealed no significant effect of these head roll position on the horizontal VOR (*F*(2.23, 24.53) = 0.16, *p* = 0.874, *η²p* = 0.02, Greenhouse-Geisser corrected).

**Figure 6.**
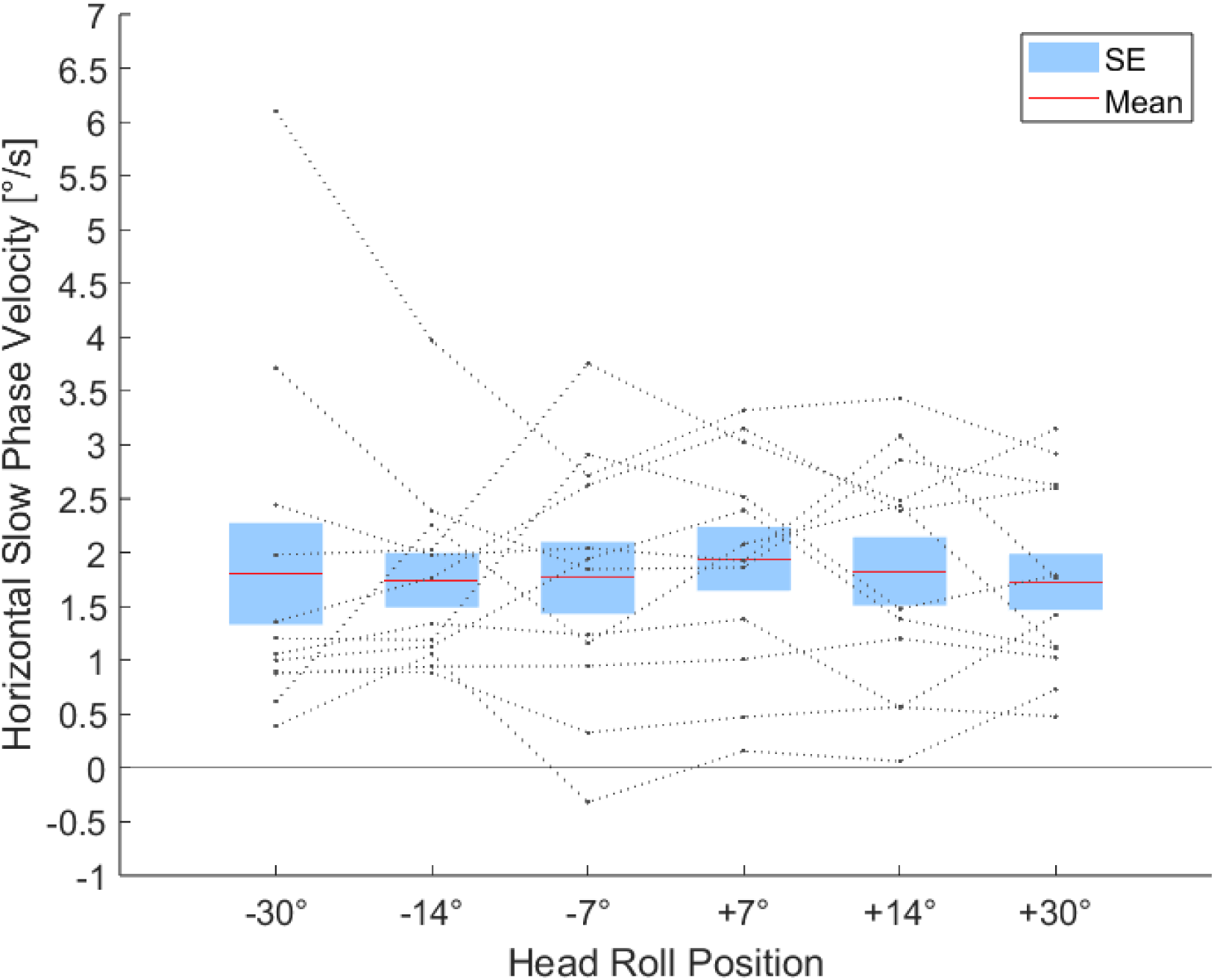
Horizontal slow phase velocity [°/s] for head roll positions across all subjects. Dotted lines represent individual subject data (circles, N = 12) for the six tilted head positions. Positive values indicate rightward slow phase, negative values indicate leftward direction. The red lines and blue boxes reflect the mean slow phase velocity and standard error (SE) across all subjects. Head roll position 0° is not depicted for simplicity, as this condition was always measured first (as a manipulation check) while all other positions were probed in a counterbalanced order and therefore showed an increased velocity due to habituation.

### 3.3 Experiment 3: Influence of combined head pitch and head roll position inside a 3 T MRI scanner

The linear regression analysis results obtained from the first two experiments (see above) revealed the average intercepts at which horizontal and vertical MVS effects were nulled. This combined 25° pitch and 15° roll head position (rounded values) is termed “MVS_null_-position” in the following. This position was contrasted with the usual supine lying position (0° pitch, 0° roll), termed “MVS_standard_-position”, and the backward tilt (-30° pitch, 0° roll) position, termed “MVS_max_-position”, in which we expected to observe the strongest impact of MVS (cf. Experiment 1). The distribution of individual horizontal and vertical slow phase velocities is shown in Figure 7. In the MVS_max_-position, participants exhibited a mean horizontal slow phase of 3.32°/s (SE= 0.79°/s) and a vertical velocity of 0.29°/s (SE= 0.39°/s). Bayesian analysis supported the presence of a horizontal VOR in this condition (BF = 562.69), while the evidence for vertical VOR was inconclusive (BF = 0.62). In the MVS_standard_-position, horizontal slow phase velocity averaged 2.08°/s (SE = 0.59°/s) and vertical –0.30°/s (SE = 0.38°/s), with strong evidence again for a horizontal VOR (BF = 83.01), and inconclusive evidence for the vertical component (BF = 0.63). In contrast, in the MVS_null_-position, velocity values were markedly reduced: horizontal slow phase averaged 0.22°/s (SE = 0.31°/s), and vertical 0.07°/s (SE = 0.20°/s). Bayes Factors in this condition favored the null hypothesis for both horizontal (BF = 0.24) and vertical (BF = 0.28) components, indicating moderate evidence for the absence of a VOR at the combined 25° pitch and 15° roll head position.

**Figure 7.**
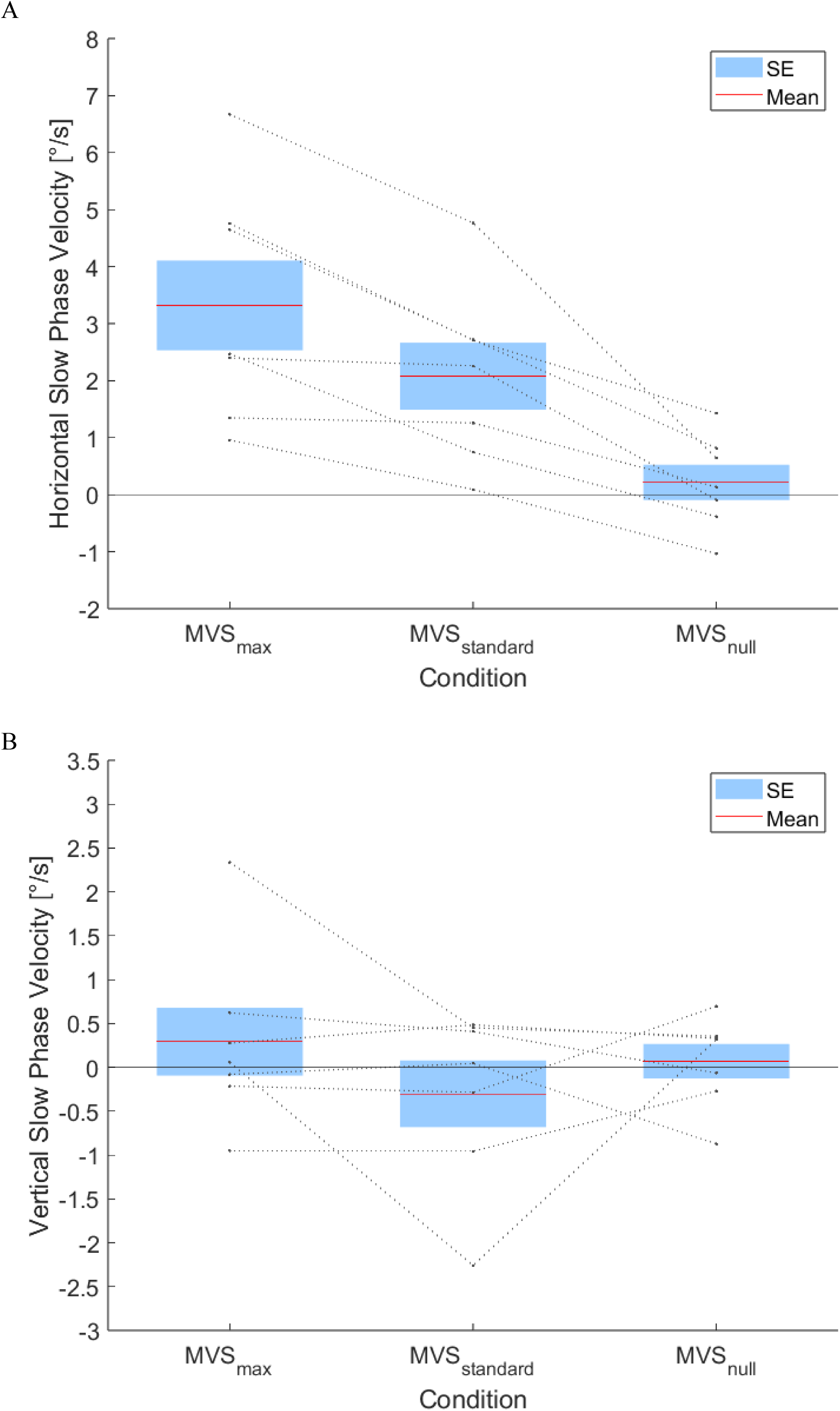
Slow phase velocity [°/s] for head positions across all subjects. Dotted lines represent individual subject data (circles, N = 7) for the three head positions. (A) horizontal VOR. (B) vertical VOR. Positive velocity values indicate rightward respectively upward slow phase movements, negative values indicate leftward respectively downward slow phase movements. The red lines and blue boxes reflect the mean slow phase velocity and SE across all subjects.

## 4. Discussion

We investigated how head position within a 3 T MRI scanner (Siemens Prisma) influences the VOR, specifically whether varying head pitch and roll can nullify MVS-induced vestibular stimulation. In Experiment 1 we confirmed that the horizontal VOR is modulated by head pitch, with a null point at a median head angle of 24.4° towards the chest. However, a consistent downward vertical VOR was observed across all pitch positions and remained unaffected. In Experiment 2, we showed that this vertical component can be modulated by head roll, with the vertical VOR nullified at an average roll of 15.9° toward the right shoulder.

It has been demonstrated in previous studies that head pitch can modulate the MVS-induced horizontal VOR responses at 7 T and that the resulting slow phase velocity is proportional to the strength of the static magnetic field (3 T vs. 7 T), although the angle at which nulling occurs appeared to vary considerably between individuals (Roberts et al., 2011). Accordingly, subsequent research showed for 3 T and 1.5 T that the MVS-induced horizontal VOR can be reduced, nulled, or even reversed when the head is tilted forward towards the chest (Boegle et al., 2016). Changes in vertical VOR with head roll have also been observed, including the possibility of reversing the direction of vertical slow phase by tilting the head toward either shoulder (Roberts et al., 2011). Based on individual MRI-based anatomical measurements of anatomical alignment of the vestibular organs relative to the magnetic field vector, an average rightward head roll of approximately 5-8° had been estimated to null the vertical VOR (Go et al., 2022), which is notably smaller than the 15.9° head roll at which we empirically observed vertical VOR-nulling.

In contrast to these prior studies, in the present Experiments 1 and 2, we measured the VOR as a function of the experimentally varied head position to individually calculate the specific pitch angle at which the horizontal VOR in a given participant was nullified and the roll angle at which an individual’s vertical VOR was extinguished, respectively. This approach further allowed us to identify an average head pitch and roll orientation that should minimize MVS-induced horizontal and vertical slow VOR components. In the subsequent Experiment 3, when we combined these head pitch and roll positions at which MVS-induced eye movements were nullified in Experiments 1 and 2, by tilting participants’ heads 25° forward and 15° to the right (rounded values), our slow phase velocity data provided evidence for the absence of both horizontal and vertical VOR components.

To explain such dependencies between head position and eye movements, Arán-Tapia et al. (2025) recently developed a biophysical model, which predicts slow phase velocity based on head orientation relative to the b_0_-vector of a static magnetic field. In a strong magnetic field, these forces interact with endogenous ionic currents, primarily flowing from utricular dark cells toward the hair cells, causing endolymph movement and cupula deflection. The resulting VOR is determined by the integration of signals from all semicircular canals. For horizontal eye movements, this integration results from the opposing activity of the left and right horizontal cristae. For instance, when the right horizontal cupula is inhibited and the left is excited, the model predicts a leftward eye movement proportional to field strength. Vertical eye movements arise from the combined activity of the superior and posterior cristae on both sides, whose opposing effects can cancel each other out. The model predicts that the horizontal VOR is nullified at a pitch angle of 27° toward the chest, which closely aligns with the 24.4° null point observed in our experimental data. For the vertical domain, the model predicted the vertical VOR to be zero at a 0° roll angle. The authors provide preliminary data from 5 subjects supporting this prediction. The difference to our vertical estimate is likely because Arán-Tapia and colleagues used vertical slow phase velocity values that were normalized to a measurement outside the MRI scanner. As this outside measurement was also obtained in the supine position, however, they thereby removed any MRI-independent “supine effect” on the vertical VOR (Bisdorff et al., 2000; also compare introduction). In contrast, our non-normalized eye data confirm that, across subjects, a vertical VOR is present at 0° head roll angle and that its magnitude can be modulated by MVS through changes in head roll position. Consequently, the combined MVS and supine effect therefore led to a zero vertical VOR when the head roll angle amounts to 15.9° towards subjects’ right shoulder.

As further predicted by the model of Arán-Tapia et al. (2025), vertical VOR was not influenced by head pitch in our study. Moreover, the model suggests that also the horizontal VOR should be modulated by head roll, although considerably less than the vertical VOR. Yet, no modulation of the horizontal VOR was observed within the range of roll-angles tested. Instead, we only detected an adaptive change in the horizontal VOR over time, which is consistent with previous observations (Smaczny et al., 2024).

In addition to the vertical and horizontal VOR components, torsional eye movement can also be elicited by MVS. Unfortunately, torsional movements could not be monitored with our eye-tracking equipment. The model by Arán-Tapia et al. (2025) suggests that the torsional VOR can also be modulated by head pitch position. However, these adjustments simultaneously influence the horizontal component with another phase-relation. Thus, the torsional VOR can only be nullified at another head pitch angle than that needed for nulling the horizontal VOR. Consequently, according to this model, no head position exists in which all three VOR components can be completely nullified; a residual torsional response thus is likely to remain. This implies that while MVS-induced eye movements can be minimized by tilting participants’ heads 25° forward and 15° to the right (shown in the present study), they very likely cannot be eliminated entirely (due to a possibly remaining torsional component).

Nevertheless, the results of our study are highly relevant for all future fMRI experiments. They demonstrate that the previously unavoidable mixture of neuronal activation in the (cortical) projection areas of the vestibular system provoked by the signal of interest and by the signal elicited by activation of the vestibular system − when participants undergo fMRI in the conventional supine position − can be significantly reduced. By tilting participants’ heads 25° forward and 15° to the right, both horizontal and vertical VOR components are nullified. Importantly, the absence of the VOR in this combined tilt condition could not be explained by VOR habituation. This is because the respective measurement was always performed in between the standard supine and the backward tilt condition, in both of which an MVS-induced VOR was detectable. Beyond reducing MVS-induced changes in neural activity, this combined head 25° forward and 15° to the right position might have further advantages, such as reducing MVS-induced shifts in oculomotor behavior or spatial attention (e.g. Lindner et al., 2021). This approach may also have clinical relevance, as exposure to strong magnetic fields can induce a sensation of body rotation, leading to dizziness or discomfort in patients (Mian et al., 2016). Optimizing head orientation to minimize MVS-induced effects could therefore contribute to improved patient comfort and overall well-being during MRI procedures.

However, achieving the optimal head position to minimize MVS-induced vestibular responses remains challenging in both basic research and clinical practice. While some newer MRI systems offer head coils with adjustable pitch mechanisms, these are not yet standard across all clinical or experimental settings. Moreover, the degree of tilt required to reach the ideal 25° pitch position is extreme and, to the best of our knowledge, not yet implemented by MRI manufacturers. For more slender individuals with smaller heads, such adjustments may be feasible within these specialized coils, but for more corpulent patients or individuals with larger head volume, attaining the necessary position becomes nearly impossible due to anatomical and spatial constraints. A further complication for basic research using visual stimulus material could result from the altered gaze angles introduced by the combined 25° pitch and 15° roll head position: Subjects might no longer be able to view visual stimulation devices via mirrors and the use of eye-trackers might no longer be viable.

In addition to these practical limitations, individual anatomical variability must also be considered. Asymmetries in the vestibular labyrinth and differences in semicircular canal orientation may lead to residual VOR even when the empirically derived average pitch and roll angles are applied. Nevertheless, the identified average position is likely to attenuate MVS-induced eye movements in most individuals. Even moderate adjustments within tolerable limits may reduce vestibular side effects and contribute to (i) a “purer” signal in neuroscientific fMRI experiments, one that is less affected by the neural activation provoked by stimulation of the vestibular system, and (ii) greater patient well-being during clinical MRI procedures.

