## Supplemental Figures for "Minimizing the influence of magnetic vestibular stimulation inside MRI-scanners by adjusting head position"

### Supplementary Figure 1: Exemplary participant from Experiment 1.

The figure presents the eye movement data for each head position from one representative participant in Experiment 1. The red lines indicate the median slow phase velocity of the VOR, where positive values represent a rightward-directed slow phase and negative values indicate a leftward direction. The horizontal VOR (Figure S1, first column) varied across head pitch position. When the head was pitched -30° backwards, the participant exhibited a rightward horizontal VOR, which progressively decreased with increasing head pitch. At the standard position with 0° pitch, the horizontal slow phase velocity was reduced compared to -30°, and with a forward pitch of +30°the direction reversed, resulting in a leftward horizontal VOR. In contrast, the vertical VOR remained consistently downward directed across all three head pitch positions (Figure S1, second column). This participant's data reflects the overall trends observed in the responses to the MVS across all 10 subjects.

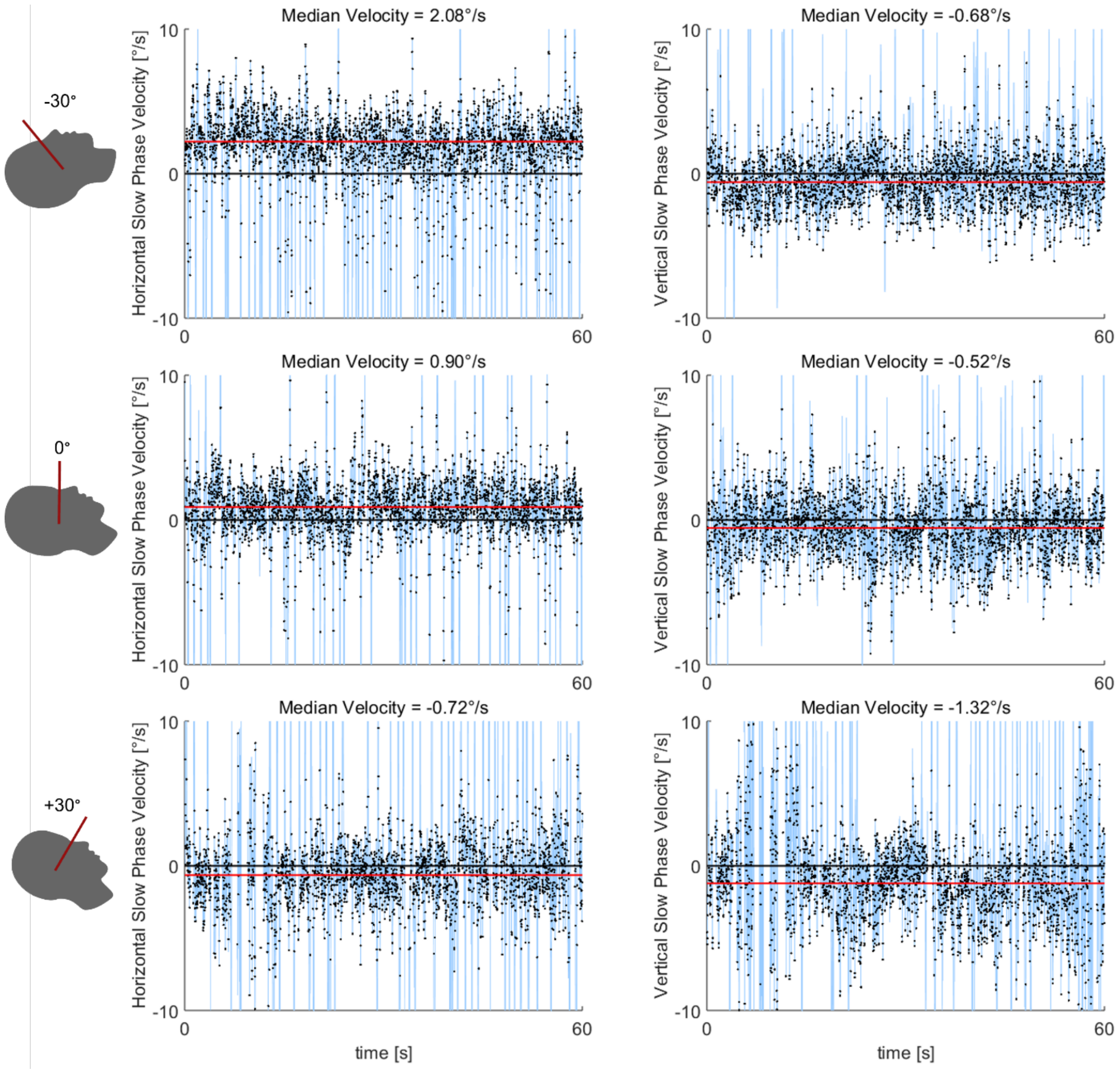


**Figure S1.** Exemplary data from a single subject showing horizontal and vertical eye velocities [°/s] of the VOR during the one-minute Nystagmus assessment. First column depicts the horizontal VOR: positive velocities represent rightward movements, while negative velocities indicate leftward movements. Seconds column shows the vertical VOR: positive velocities correspond to upward movements, and negative velocities to downward movements. Each row corresponds to one of the three head pitch positions (-30°, 0, 30°). Blue vertical lines mark saccades, while black dots indicate saccade endpoints. The red horizontal line represents the median horizontal and vertical slow phase velocities.

**Supplementary Figure 2: Exemplary participant from Experiment 2.**

The figure presents the eye movement data across head position from one representative participant in Experiment 2. Red lines indicate the median slow phase velocity of the VOR, with positive values denoting rightward directed slow phase and negative values indicating leftward directions. The horizontal VOR (Figure S2, first column) remained constantly rightward directed across head roll position. In contrast, the vertical VOR (Figure S2, first column) varied with head tilt. When the head was tilted -30° to the left, a downward directed slow phase was observed. As the head was tilted progressively to the right, the vertical VOR decreased and eventually reversed direction, becoming upward directed at 14° rightward roll. This participant's data exemplifies the overall trends observed in response to MVS across all 12 participants.

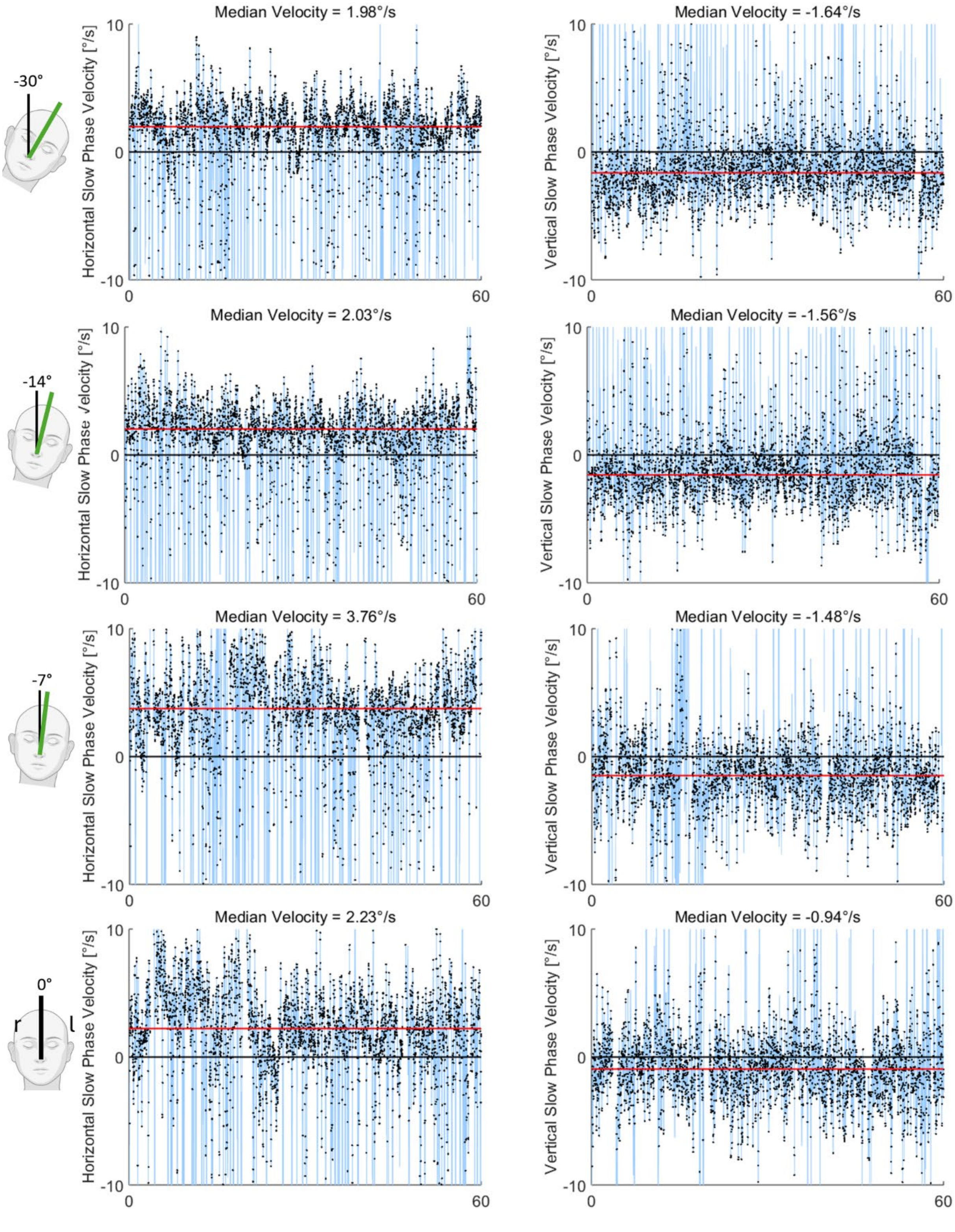
-30°
-14°
-7°
0°
r
l
Median Velocity = 1.98°/s
Median Velocity = -1.64°/s
Median Velocity = 2.03°/s
Median Velocity = -1.56°/s
Median Velocity = 3.76°/s
Median Velocity = -1.48°/s
Median Velocity = 2.23°/s
Median Velocity = -0.94°/s
Horizontal Slow Phase Velocity [°/s]
Vertical Slow Phase Velocity [°/s]
10
0
-10
60

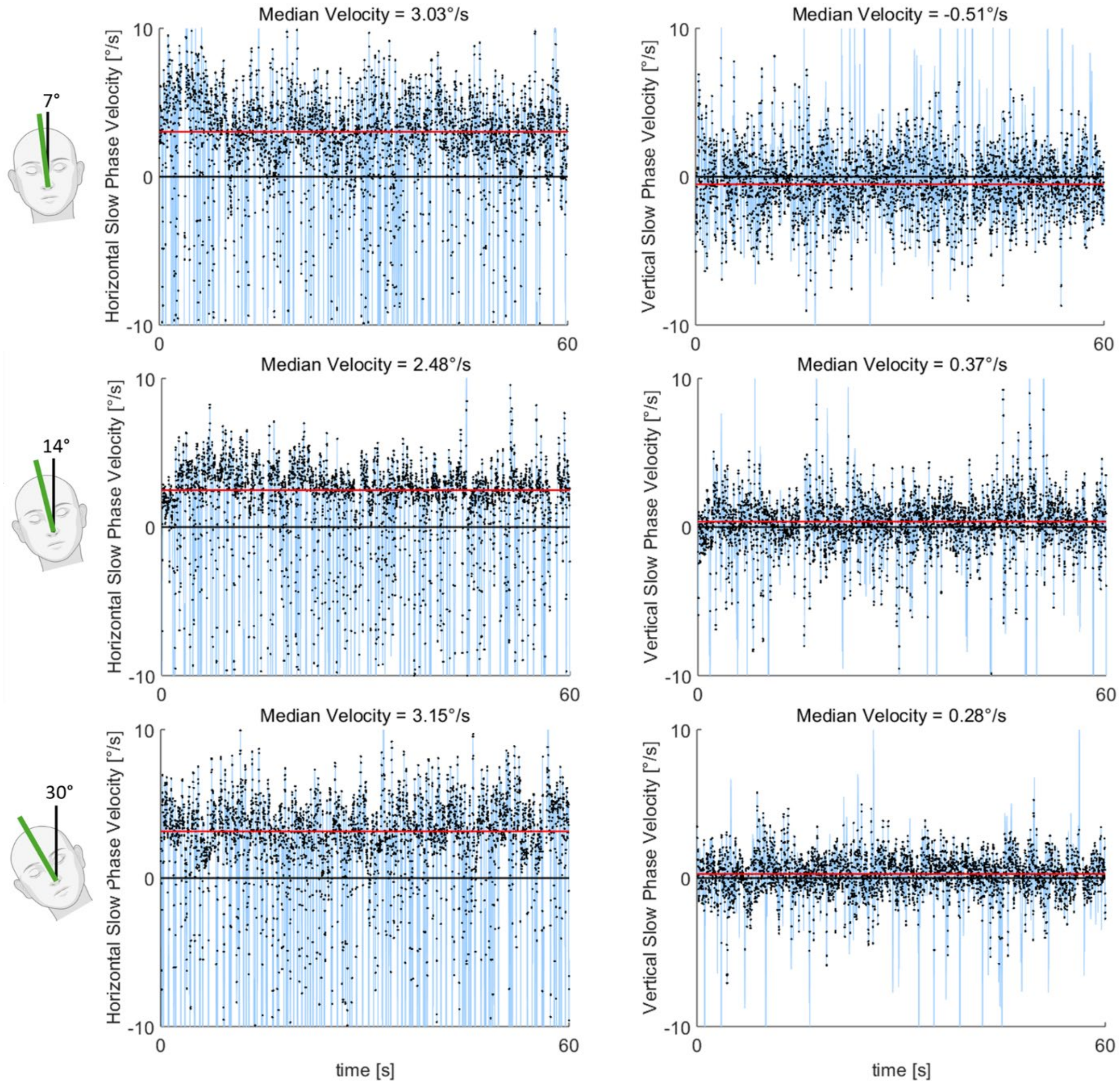


**Figure S2.** Exemplary data from a single subject showing horizontal and vertical eye velocities [°/s] of the VOR during the one-minute Nystagmus assessment. First column depicts the horizontal VOR: positive velocities represent rightward movements, while negative velocities indicate leftward movements. Seconds column shows the vertical VOR: positive velocities correspond to upward movements, and negative velocities to downward movements. Each row corresponds to one of the seven head roll positions (-30°, -14°, -7°, 0, 7°, 14°, 30°). Blue vertical lines mark saccades, while black dots indicate saccade endpoints. The red horizontal line represents the median horizontal and vertical slow phase velocities.
